# Micro-offline gains do not drive implicit motor sequence learning

**DOI:** 10.64898/2026.08.17.745334

**Authors:** Tharan Suresh, Michael V Freedberg, Sara J Hussain

## Abstract

Motor sequence performance improves during and between brief practice bouts (micro-online and offline gains). We compared both metrics across two groups: one exposed to an implicit motor sequence, and one not. Micro-online gains drove sequence-specific learning and positively correlated with overall skill. However, micro-offline gains were comparable between groups and did not track sequence-specific learning. We conclude that implicit motor sequence learning is driven by micro-online rather than micro-offline gains.

## Main text

The ability to plan, execute, and learn sequential actions is critical for everyday life^1^. Sequence execution changes both during practice (micro-online gains) and across brief rest intervals between practice bouts (micro-offline gains)^1,2^, raising the possibility that both processes support learning. Recent studies using explicit motor sequence learning tasks reported that early execution improvements were driven solely by micro-offline gains, with virtually no micro-online gains^3,4^. Consequently, micro-offline gains have recently been treated as a general property of motor sequence learning and a behavioral signature of rapid, replay-mediated consolidation^5–9^. However, whether micro-offline gains drive implicit learning remains unresolved.

To date, no studies have investigated micro-offline gains during implicit sequence learning tasks using a control group where no sequence order can be learned. Without such a group, it is difficult to fully dissociate improvements in motor execution from true sequence-specific learning. Here, we resolve this fundamental limitation by comparing micro-online and micro-offline gains between two groups of healthy adult participants: one that performed the implicit serial reaction time task (SRTT) with a hidden, embedded sequence, and one that performed the random serial reaction time task (RRTT). We reasoned that if micro-offline gains drive implicit motor sequence learning, then these gains should be significantly larger in participants who performed the SRTT than those who performed the RRTT. Further, micro-offline gains should correlate with conventional sequence-specific acquisition and retention metrics.

Forty right-handed, healthy young adults (25% male) were randomly assigned to either the SEQUENCE (N=20, 14 female, *M* = 20.05, *SD* = 1.46) or NO-SEQUENCE group (N=20, *M* = 21.7, *SD* = 3.81). Participants were instructed to respond as quickly and accurately as possible to the visual cues by pressing the corresponding key using the first four fingers of their right hand (Figure 1). SEQUENCE participants performed the task with an embedded 12-item sequence (SRTT) while NO-SEQUENCE participants performed pseudorandom keypresses (RRTT). For SEQUENCE participants, acquisition blocks 1, 2, 8, and 14 were random blocks, while acquisition blocks 3-7 and 9-13 were sequence blocks. For these same participants, retention blocks 1 and 7 were random, while retention blocks 2-6 were sequence blocks. For NO-SEQUENCE participants, all acquisition and retention blocks were random. For all participants, each block contained 10 trials (120 keypresses) and consecutive blocks were separated by 60 seconds of rest. For the SEQUENCE group, acquisition skill was defined as the difference in mean RT between the last random and last sequence blocks during acquisition, and between the first random and first sequence blocks during retention. For the NO-SEQUENCE group, the difference between mean RT during the first and last block was calculated for both acquisition and retention. As per Bönstrup and colleagues^3,4^, speed was calculated as the number of correct responses per second (keypresses/s), micro-offline gains were calculated as the speed difference between the last correct sequence of a given block and the first correct sequence of the next block, and micro-online gains were calculated as the speed difference between the first correct sequence of a given block and the last correct sequence of the same block. Summed micro-offline and micro-online gains were then computed as the sum of each gain type across sequence blocks (blocks 3-7 and 9-13 for each group). Summed total gains were computed as the sum of micro-offline and micro-online gains. After retention testing, explicit sequence awareness was evaluated using the process dissociation procedure (PDP^10^).

**Figure 1.**
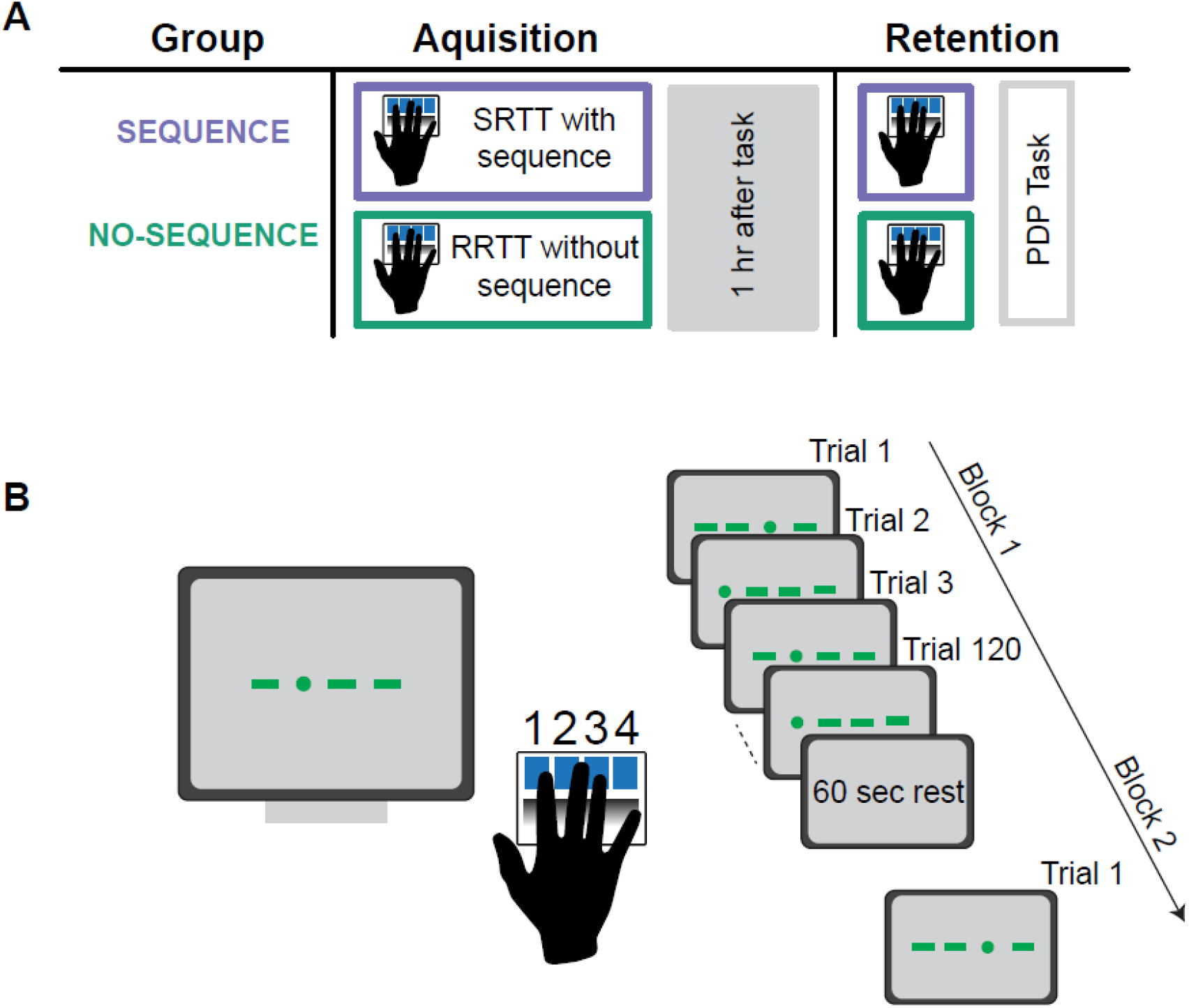
Experimental design. (A) Participants in the sequence group (SEQUENCE) performed the implicit SRTT with an embedded 12-item repeating sequence, while those in the no-sequence group (NO-SEQUENCE) performed the RRTT with pseudorandom keypresses. (B) Participants responded to a circle that appeared at one of four screen positions by pressing the corresponding key using fingers 1-4 of their right hand. Participants were not informed of the presence or absence of a repeating sequence in either group. A correct response was required to progress to the next trial. Acquisition contained 14 blocks of 120 keypresses (10 sequence repetitions per block), separated by 60 s of rest. One hour later, participants completed a retention test (7 blocks) followed by the process dissociation procedure (PDP) to test explicit sequence awareness.

During SRTT acquisition, both groups improved task performance (Figure 2A, RT: *F*_(13, 481)_ = 18.17, *p* < .001; Figure 2C, speed: *F*_(13, 481)_ = 10.36, *p* < .001). Performance did not differ between groups for random blocks, but the SEQUENCE group performed better during sequence blocks than the NO-SEQUENCE group (RT: Group × Block: *F*_(13, 481)_ = 13.97, *p* < .001; speed: Group × Block: *F*_(13, 481)_ = 9.19, *p* < .001). Post hoc comparisons during random blocks showed no group differences (t < −1.39, *p*_*adj*_ > .5, for all comparisons). By the final sequence block, the SEQUENCE group was significantly faster than the NO-SEQUENCE group (*t* > 4.3, *p*_*adj*_ < .0001 for both RT and speed).

**Figure 2.**
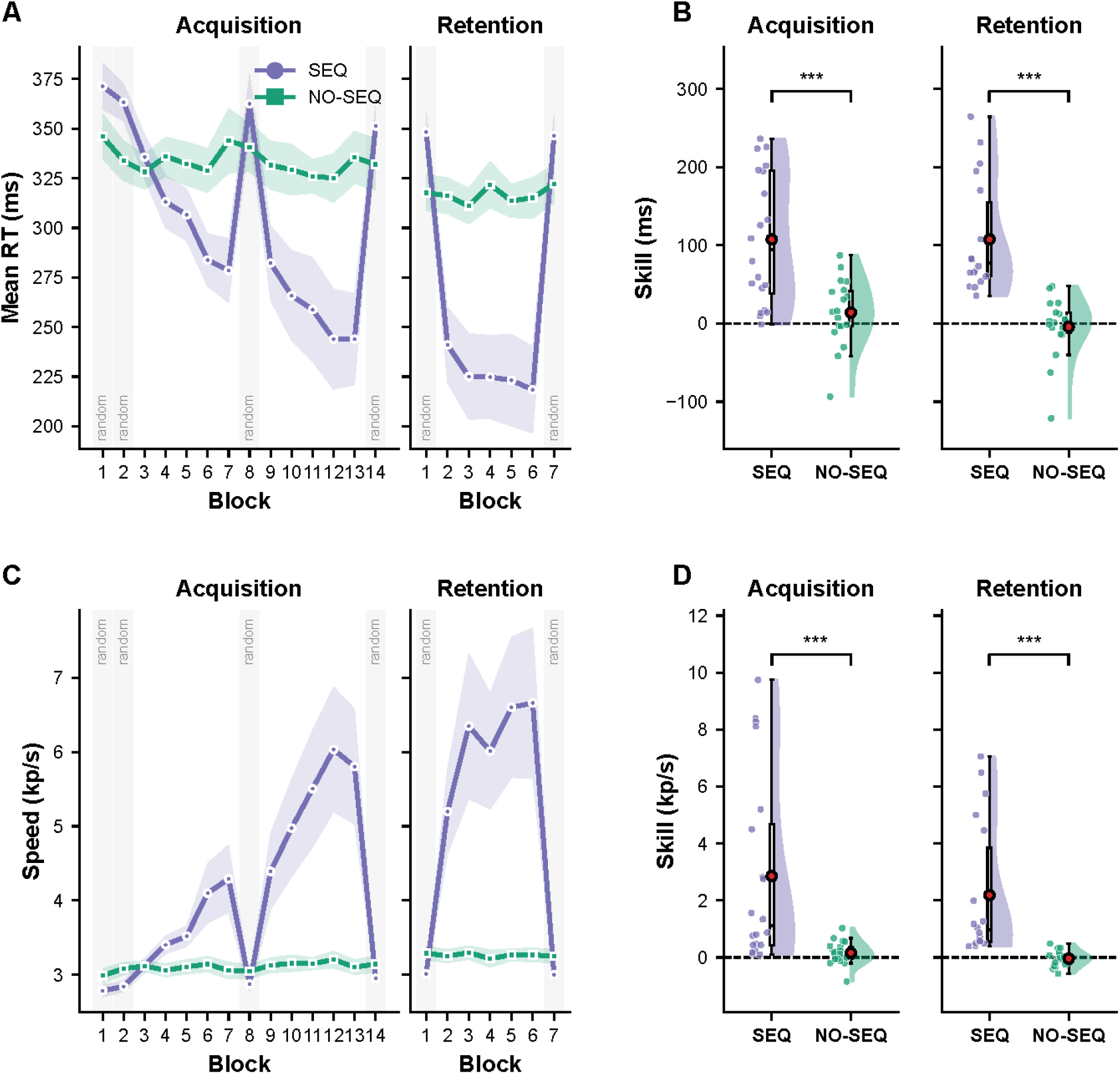
Sequence-specific skill is acquired and retained in the SEQUENCE group. (A) Mean RT per acquisition (left) and retention (right) blocks. Grey shading denotes random blocks; for the SEQUENCE group, all other blocks were sequence blocks. (B) Acquisition and retention skill (ms). (C) Speed (keypresses/s) per block, plotted as in A. (D) Acquisition and retention speed (keypresses/s). In A and C, lines and shading represent mean ± SEM. In B and D, dots represent individual participants, red circles and error bars are group mean ± SEM, boxplots show median and interquartile range, violin plots show data distribution, and dashed lines mark zero. ***p < .001; Welch’s t-tests.

A similar pattern was observed during retention: performance improved across blocks (Figure 2A, RT: *F*_(6, 204)_ = 34.52, *p* < 0.001; Figure 2C, speed: *F*_(6,204)_ = 12.83, *p* < .001), but the SEQUENCE group showed better performance than the NO-SEQUENCE group during sequence blocks (*t* > 3.53, *p*_*adj*_ < .02, for all comparisons). The SEQUENCE group also showed greater acquisition skill than the NO-SEQUENCE group (Figure 2B, Acquisition skill: *t*_(28.5)_ = 4.42, *p* < .001, *d* = 1.39; Figure 2D, Speed: *t*_(19.6)_ = 3.63, *p* = .001, *d* = 1.13) and retained this skill 1 hour later (Retention skill: *t*_(26.5)_ = 5.81, *p* < .001, *d* = 1.94; Speed: *t*_(17.5)_ = 3.98, *p* < .001, *d* = 1.33). We found no strong evidence of explicit sequence awareness in either group (SEQUENCE group inclusion test accuracy: 6.17 ± 0.67, *t*_(16)_ = 3.21, *p* = .005; SEQUENCE group exclusion test accuracy: 7.94 ± 0.52, *t*_(16)_ = −0.11, *p* = .91; NO-SEQUENCE group inclusion test accuracy: 3.88 ± 0.45, *t*_(16)_ = −0.25, *p* = .79; NO-SEQUENCE group exclusion test accuracy: 7.7 ± 0.54, *t*_(16)_ = −0.53, *p* = .59; see Figure S2).

During acquisition, summed total gains and summed micro-online gains were greater in the SEQUENCE than the NO-SEQUENCE group (Figure 3B, micro-online: *t*_(20.60)_ = 2.89, *p* = .008, *d* = 0.9; total: *t*_(21.53)_ = 4.28, *p* < .001, *d* = 1.34). However, summed micro-offline gains did not differ between groups (micro-offline: *t*_(22.1)_ = −2.00, *p* = .06, *d* = −0.63).

**Figure 3.**
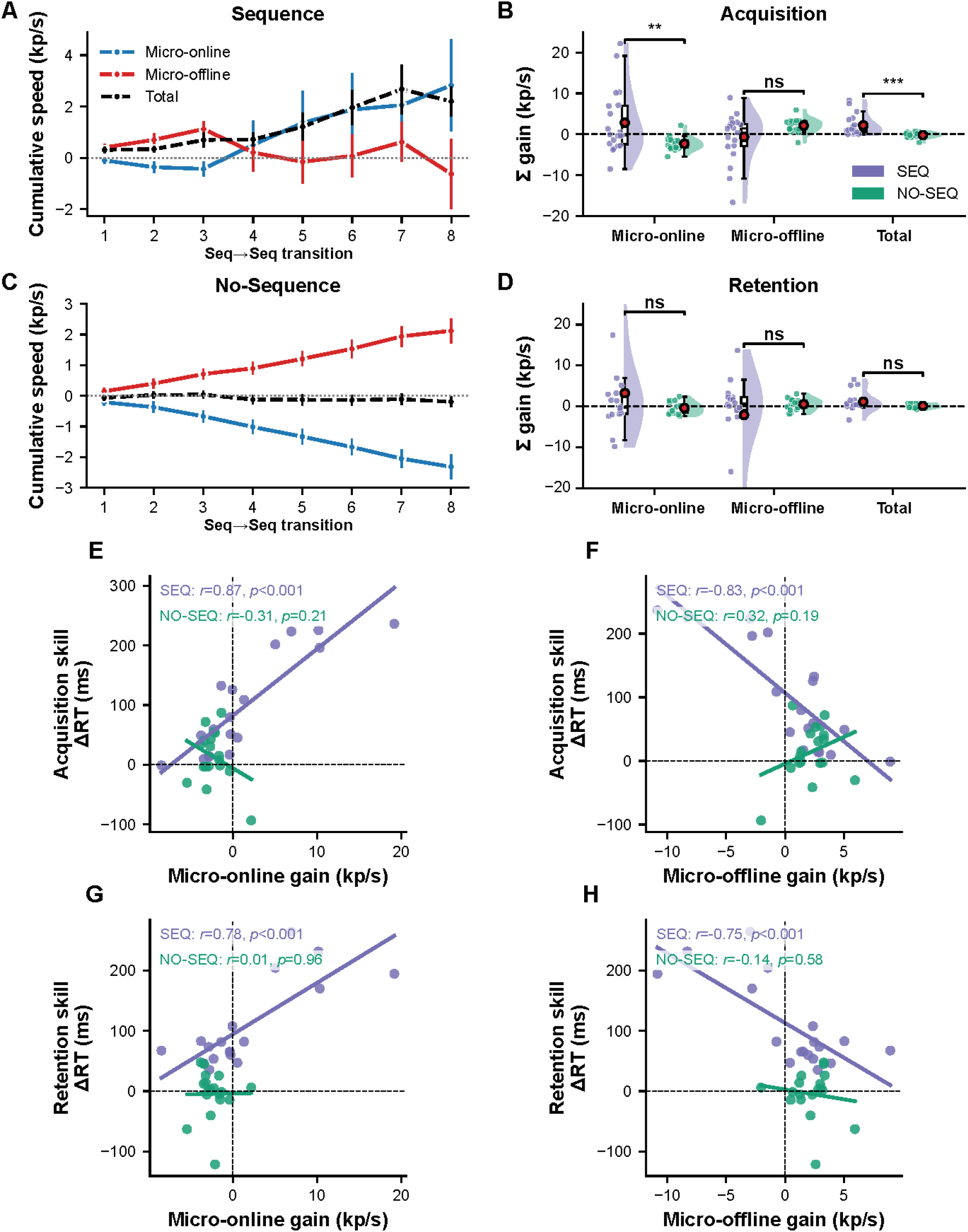
Micro-offline gains do not drive implicit motor sequence learning. (A, C) Cumulative micro-online, micro-offline, and total speed gains across the eight sequence-to-sequence block transitions during acquisition for the SEQUENCE (A) and NO-SEQUENCE (C) groups. Data are group means ± SEM. (B, D) Total micro-online, total micro-offline, and summed total gains during acquisition (B) and retention (D), plotted as in Fig. 2B. (E-H) Relationships between summed total gains and skill per group. Dots represent individual participants; lines are least-squares fits. **p < .01, ***p < .001; ns, not significant.

During retention, neither summed total, micro-online, or micro-offline gains differed between groups (Figure 3D, micro-online: *t*_(17.4)_ = 1.20, *p* = .25, *d* = 0.4; micro-offline: *t*_(17.4)_ = −0.86, *p* = .4, *d* = −0.29; total: *t*_(18.3)_ = 1.81, *p* = .08, *d* = 0.6).

In the SEQUENCE group, acquisition skill positively correlated with summed micro-online gains (Figure 3E, *r*_(16)_ = 0.87, *p* < .001) and negatively correlated with summed micro-offline gains (Figure 3F, *r*_(16)_ = −0.83, *p* < .001). This was not the case for the NO-SEQUENCE group (micro-online: *r*_(16)_ = −0.3, *p* = .21; micro-offline: *r*_(16)_ = 0.32, *p* = .19). We also evaluated whether micro-online and micro-offline gains calculated during acquisition predicted retention skill. For the SEQUENCE group, micro-online gains positively correlated with retention skill, while micro-offline gains negatively correlated with retention skill (Figure 3G, micro-online: *r*_(16)_ = 0.78, *p* < .001; Figure 3H, micro-offline: *r*_(16)_ = −0.75, *p* < .001). However, this was not the case for the NO-SEQUENCE group (micro-online: *r*_(16)_ = .01, *p* = .97; micro-offline: *r*_(16)_ = −0.13, *p* = .59).

In summary, the SEQUENCE group showed greater implicit sequence learning than the NO-SEQUENCE group. In the SEQUENCE group, micro-online gains closely tracked overall performance improvements (see Figure 3A), and micro-online gains calculated during acquisition positively correlated with both acquisition and retention skill. However, micro-offline gains did not differ between groups, and for the SEQUENCE group, micro-offline gains calculated during acquisition negatively correlated with acquisition and retention skill. These findings show that implicit sequence learning is largely driven by micro-online gains and that micro-offline gains occur even when participants are not exposed to a repeating sequence (see Figure 3C).

Why might micro-online gains drive implicit but not explicit sequence learning? First, although both tasks involve the acquisition of sequential knowledge, they recruit different cognitive processes^11^ and rely on dissimilar sensorimotor networks^12,13^. Explicit learning is driven by top-down factors like attention and effort; it also leverages short-term memory to consciously retrieve and improve sequence performance^14–18^. In contrast, implicit learning relies on bottom-up stimulus-driven processes, develops gradually through incremental exposure to stimulus-response associations, and requires fewer attentional resources^19,20^. Consistent with these differences, the implicit SRTT recruits corticostriatal circuits^12,21^, while explicit sequence learning more heavily engages prefrontal and premotor areas^22,23^. Second, explicit sequence learning is self-paced and uses short (typically 5-item) sequences that can be preplanned and executed quickly^24^. Because participants are instructed to execute sequences as fast as possible, they often experience fatigue^25,26^ and reactive inhibition^27^ that then dissipates across rest breaks. However, the implicit SRTT is externally paced, such that each element within a longer (typically 12-item) sequence is presented independently^11,28^. This design limits the accumulation of fatigue and reactive inhibition during task performance and thus results in smaller micro-offline gains that do not track performance improvements. Third, performance gains during the implicit SRTT emerge slowly over many repetitions^11,28^. As a result, performance curves during this task lack the characteristic sharp initial rise and subsequent performance plateau observed during explicit sequence learning^29,30^. Because performance gains during the implicit SRTT accrue across the entire learning session^29^, we calculated micro-offline and micro-online gains across all sequence-to-sequence acquisition block transitions. This analytical difference may explain why prior implicit SRTT studies observed minimal micro-online gains^31^. However, by analyzing the full acquisition period and including a NO-SEQUENCE control group, our study experimentally isolates the role of micro-online gains in implicit sequence learning.

Micro-offline gains conflate at least four processes: genuine skill learning, motor preparation^32,33^, recovery from peripheral fatigue^25,26^, and dissipation of reactive inhibition^27^. Our experimental design helps untangle these contributions. First, speed declined during each block and recovered after rest in both groups (see Figure S1); these declines were likely driven by fatigue and reactive inhibition. However, in the SEQUENCE group, within-block speed declines were offset by concurrent sequence-specific learning. Because the NO-SEQUENCE group involves identical motor execution and rest demands, it directly quantifies non-sequence-specific speed changes. Under these well-controlled conditions, micro-offline gains did not differ between groups. These findings align with recent work suggesting that micro-offline gains reflect dissipation of reactive inhibition^27^ rather than sequence-specific learning or rapid consolidation^32,34,35^. Second, because the implicit SRTT cues one keypress at a time and uses a relatively long sequence, the contribution of sequence pre-planning to micro-offline gains is lower than in explicit sequence learning tasks^36^. Consistent with this, attenuating pre-planning during explicit sequence learning sharply reduced micro-offline gains^32^, and incorporating motor preparation time into micro-offline metrics flips gains into losses^33^. Third, prior studies have shown that inter-block rest duration does not alter total learning^37,38^ but instead restructures when performance improvements occur. That is, shorter and longer rests favor micro-online and micro-offline improvements, respectively^37^. In our study, both groups experienced identical 60-second rest breaks that were likely long enough to recover from any fatigue and reactive inhibition that accumulated during execution periods. Consequently, micro-offline gains were comparable across groups, suggesting that they reflect task-general processes rather than sequence-specific learning or rapid consolidation.

Overall, our findings indicate that implicit sequence-specific learning is driven by micro-online but not micro-offline gains. These results suggest that micro-offline gains reflect task-general processes that do not require sequence learning. Further, experimentally isolating the relative contributions of these two metrics to overall learning is only possible when including a control group that is not exposed to any repeating sequence. Without such a group, micro-offline gains can be mistaken for sequence-specific learning.

## Methods

### Experimental design

Forty right-handed healthy adults participated in a larger experiment that included the tasks described here^39^. Right-handedness was verified using the Edinburgh Handedness Inventory. Participants were randomly assigned to either the SEQUENCE (N=20) or the NO-SEQUENCE group (N=20). Participants in the SEQUENCE group performed the implicit serial reaction time task (SRTT) with a hidden 12-item repeating sequence, while those in the NO-SEQUENCE group performed the random reaction time task (RRTT). One hour after acquisition, both groups performed a brief retention test using the same task version. The process dissociation procedure (PDP) was also used to evaluate sequence awareness. Of the 40 participants, 1 was excluded for left-handedness, 3 did not complete retention testing, and 5 did not complete the PDP. As a result, the sample size varied across analyses and is reported for each result. The study was approved by the Institutional Review Board at The University of Texas at Austin, and all participants provided written informed consent prior to participation.

### Serial and Random Reaction Time Tasks

Participants performed either the implicit SRTT or RRTT, each of which were programmed in E-Prime 3 (Psychology Software Tools). Each trial presented a circle at one of four screen positions, and participants were asked to respond to the circle by pressing the spatially compatible key with their right hand as quickly and accurately as possible (index = 1, middle = 2, ring = 3, pinky = 4). A correct response was required to advance to the next trial. The SEQUENCE group completed 14 acquisition blocks of 120 keypresses each (10 repetitions of a 12-item sequence: 2-3-1-4-3-2-4-1-3-4-2-1), with blocks 1, 2, 8, and 14 as random blocks. The NO-SEQUENCE group performed pseudorandomized keypresses during all blocks. Blocks were separated by 60 s of rest during which participants rested quietly. Both groups completed a 7-block retention test one hour after acquisition (random blocks 1 and 7; sequence blocks 2-6). Participants were not informed of the presence or absence of a repeating sequence in either group.

### Reaction time-based skill measures

Reaction times (RTs) were averaged within each block. For the sequence group, acquisition and retention skill were defined as the difference in mean RT between the last random and last sequence block during acquisition, and between the first random and first sequence block during retention, respectively. Because the NO-SEQUENCE group performed pseudorandom keypresses in all blocks, acquisition and retention skill for this group were defined as the difference in mean RT between the first and last block of the corresponding phases.

### Micro-gains analysis

Keypress speed was computed using previously established metrics^3^. Mean inter-keypress interval (IKI) per sequence was converted to keypresses/sec (1000 / mean IKI). Micro-online gains were calculated as the difference in speed between the last and first sequence within a block; micro-offline gains were calculated as the difference in speed between the first sequence of block *n+1* and the last sequence of block *n*. Only sequence-to-sequence block transitions were included in micro-gain analyses. Cumulative micro-gains were computed by calculating the running arithmetic sum across these blocks. Total micro-gains were calculated by summing gains across acquisition and retention. Acquisition and retention speed were calculated using the same approach used for RT-based metrics.

### Process Dissociation Procedure

Following retention, sequence awareness was assessed using the process dissociation procedure (PDP), which indexes conscious control over any sequence knowledge acquired during implicit SRTT performance. This procedure is based on the rationale that conscious knowledge is both reportable on request and under intentional control. The PDP comprised an inclusion and an exclusion test, each of which contained 12 unique three-item fragments from the 12-item sequence embedded in the SRTT (e.g., 2-3-1, 3-1-4, 1-4-3). The inclusion test evaluates reportable, consciously accessible knowledge. Here, participants were shown the first two elements of each fragment and asked to produce the next element. The exclusion test evaluates intentional control. Here, participants were shown the first two elements of each fragment and asked to produce any element *other* than the one that completed the fragment. Participants were instructed that immediate element repetitions (e.g., a 2 immediately after a 2) were not permitted. Consequently, chance performance was 1/3 (4 of 12) for the inclusion test and 2/3 (8 of 12) for the exclusion test. Test order was counterbalanced across participants per group. For each group, the number of accurate inclusion and exclusion trials was compared against its respective chance level using one-sample t-tests.

### Statistical Analysis

All data and statistical analyses were performed in R (version 4.5.0) and Python (version 3.13). Reaction time and speed were analyzed using linear mixed-effects models (lmerTest) with Group, Block, and their interaction as fixed effects and individual participants as random intercepts. F-tests used Type III sums of squares with degrees of freedom estimated using Satterthwaite approximation. Post hoc tests were performed when the Group × Block interaction was significant. Group differences at random probe blocks and the last sequence block were tested using estimated marginal means with Holm-Bonferroni correction, and effect sizes were derived from the model’s residual standard deviation. Between-group differences in RT skill and speed, summed micro-online, summed micro-offline, and summed total gains were each tested using independent-samples t-tests (Welch’s correction for unequal variances). Cohen’s *d* calculations were based on the pooled standard deviation. Relationships between summed micro-gains and skill were assessed per group using Pearson’s correlations. PDP accuracies were compared against chance using one-sample t-tests. Alpha was 0.05 (two-tailed) for all analyses.

## Data and code availability

The data and code used in the study are available on GitHub at https://github.com/tharan010/Micro-gains-during-implicit-sequence-learning.git

## Acknowledgements

We would like to thank Maggie McElmurry and Muskan Manesiya for their help with data collection and Nafiz Ahmed for the helpful discussions.

## Author contributions

TS collected and analyzed the data for this project. TS, MVF, and SJH designed the experiment and wrote the manuscript. SJH supervised the project. All authors read and approved the final version of the manuscript.

