## Supplementary material for "Micro-offline gains do not drive implicit motor sequence learning"

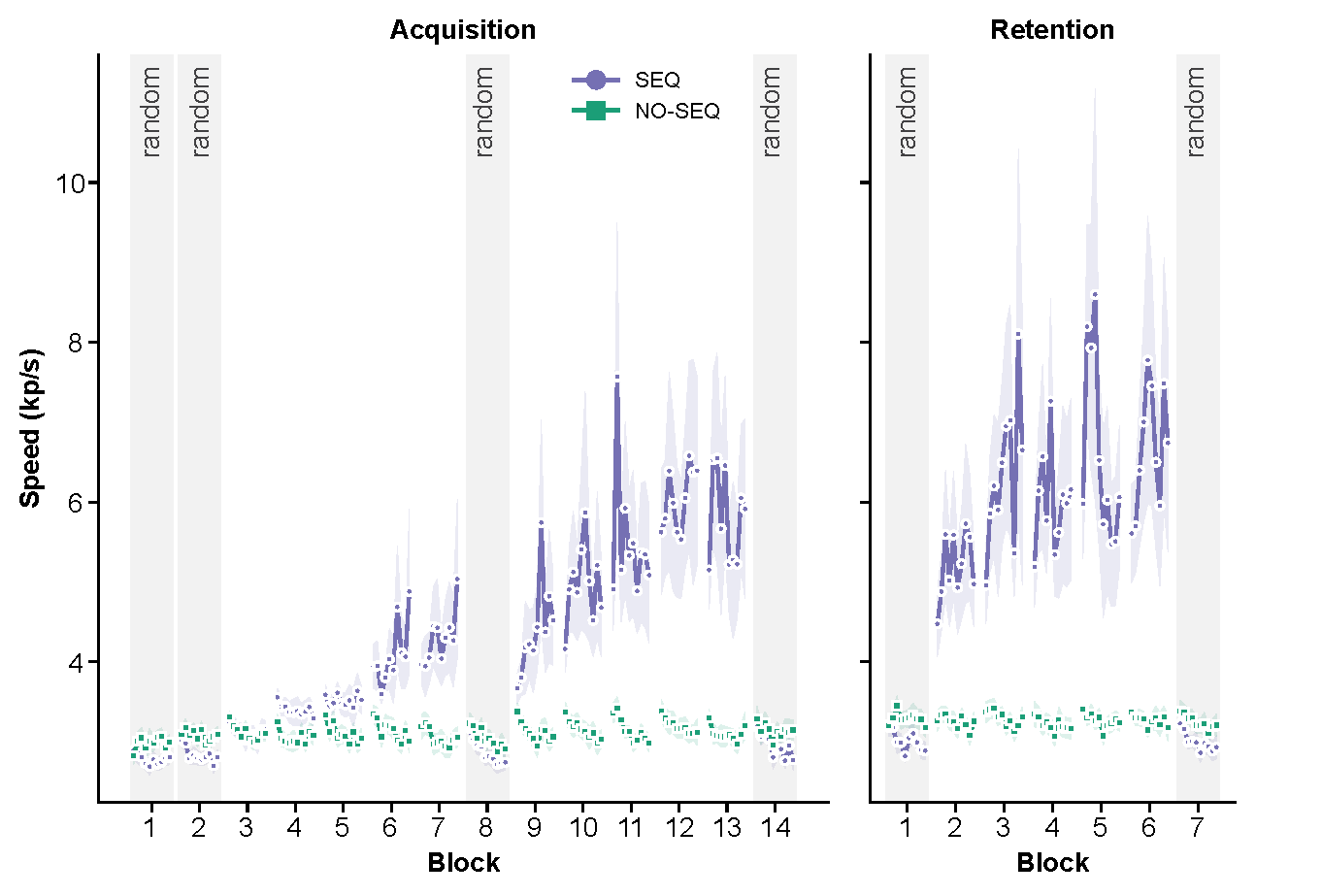
**Supplementary Figure S1.** Speed (keypresses/s) per trial within each block during acquisition (left) and retention (right) blocks. Each block contained 10 trials, with 120 keypresses each. Grey shading denotes random blocks; for the SEQUENCE group, all other blocks were sequence blocks. Dots represent individual participants, lines and shading represent mean ± SEM.

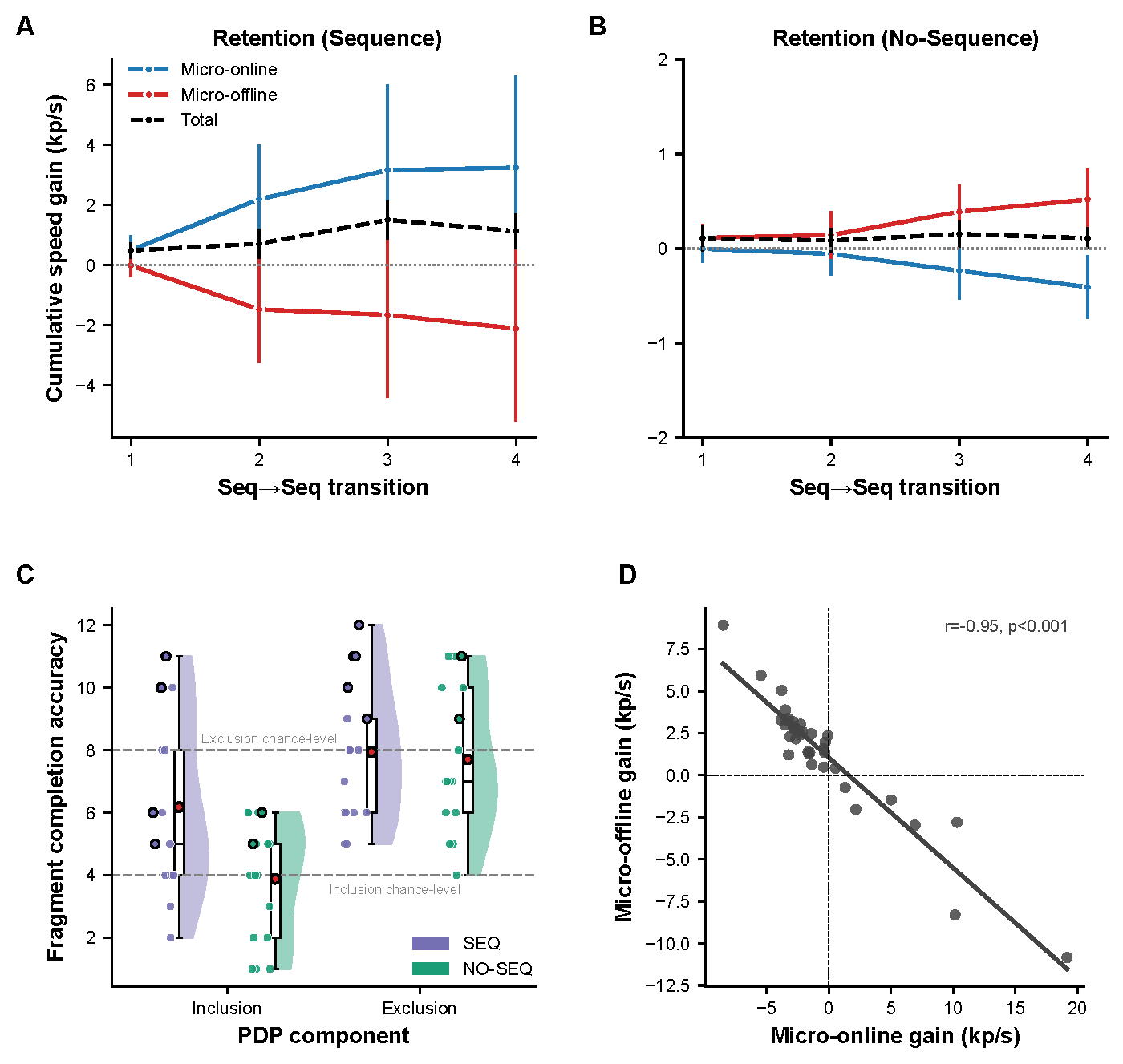
**Supplementary Figure S2.** (A, B) Cumulative micro-online, micro-offline, and total gains across the four sequence-to-sequence block transitions during retention for the SEQUENCE (A) and NO-SEQUENCE (B) groups. Data are group means ± SEM. (C) PDP fragment completion accuracy for inclusion and exclusion tests per group, plotted as in Fig. 2B. Dashed lines mark chance (4/12 for inclusion and 8/12 for exclusion). Dots with black circles represent participants who gained explicit sequence knowledge. (D) Strong negative correlation between total micro-online and micro-offline gains across all participants (r = -0.95, p < .001).
